# Evidences for newer insecticides and growth regulators resistance in field populations of *Helicoverpaarmigera* (Lepidoptera: Noctuidae) in Karnataka,India

**DOI:** 10.1101/2022.01.02.474718

**Authors:** Honnakerappa S. Ballari, Shashikant. S. Udikeri, Vinay Kalia

## Abstract

The prominence of *Helicoverpaarmigera*(Lepidoptera: Noctuidae)averse insecticide resistance was traversed in the course of 2017 in Karnataka,India. The results divulged typical resistance level prostrating in selected newer insecticides, even though exiguous higher resistance airing in insect growth regulator Novaluranwith LC 50of and 13.02 to 18.07 ppm and 1.17 to 1.95 folds resistance) compared to newer group insecticides Spinasad, Indoxacarb, Flubendiamide and Rynaxypyr(3.57 to 10.19 ppm, 1.01 to 1.27 fold). Raichur and Kalaburgi strains comprehend higher resistance to Novaluran and newer insecticides with exception of Flubendiamide (Raichur and Vijayapura strains), respectively and Spinosad (Kalaburgi and Raichur strains), respectively. The morphometric parameters of larval length, pupal length, and weight were most in RCH stain (2.75±0.48 cm, 1.76±0.18 cm, and 0.511±0.04 g, 0.309±0.05 g) respectively, which was pursued by Kalaburgi strain. The morphometric correlation revealed that larval length was a significant positive relation with insecticide resistance which might be an influence of resistance but not merely responsible. Among newer insecticides, a significant positive correlation between Rynaxypyr and Indoxacarb was evident, similarly, Nuvaluran with Indoxacarb and Rynaxypyr as well. Usage pattern revealed that 81.67 % of farmers found to use insecticides more than the recommended dose and 70.83% have habit consecutive applications of products from the same chemical group which bear witness to developing resistance.

## Introduction

*Helicoverpaarmigera* (Hubner) is a cosmopolitan insect pest that is widely distributed in the world. This pest has a broadly preferred host with a range of over 360 plant species which also includes the crop plants *viz*, cotton, maize, sorghum, sunflower, tomato, okra and legumes (Singh and Singh 1975). Pawar*et al*. (1986) found 182 plant species as hosts of *H. armigera*of which 56 were heavily damaged and 126 were rarely affected. Worldwide, losses due to *Helicoverpa* in cotton, legumes, vegetables, cereals, etc., exceed US$
2 billion and the cost of insecticides used to control these pests is over US$
1 billion annually (Reed and Pawar 1982). In India, a yield loss caused by *H. armigera in* differentcrops was in the range of 20 to 30 percent and sometimes rose to 75 percent in chickpea (Rahman, 1989) and range from 70 to 95% (Prakash*et al*., 2007). In cotton, the loss was reported in Tamil Nadu and Karnataka likely to be 35-38 % and insecticides worth 28800 billion rupees are used annually on all crops in India of which half are used on cotton alone (Rai*et al*., 2009).

*H. armigera* is high polyphagy, wide geographical range, mobility, migratory potential, facultative diapause, high fecundity (Fitt, 1989) and propensity to progress insecticides resistance are physiological, ethological and ecological factors that have robustly subsidized to its pest stature which able to endorse various cropping systems. In the 1990s the most reported cases of insecticide resistance worldwide with evolved resistance against pyrethroids, organophosphates and carbamates but recently against the *Bacillus thuringiensis* derived toxins. However, with the extensive use of chemicals, widespread resistance to insecticides crop up in *H. armigera* in India.

Pest has been wreaked to heavy selection pressure and the development of resistance to the major chemical families of insecticides has been recorded, including carbamates(methomyl, thiodicarb, carbaryl), organophosphates (monocrotophos, quinalphos and phoxim, and to a lesser extent profenofos, methyl-parathion, phosalone and chlorpyrifos) and especially pyrethroids (i.e. permethrin, fenvalerate, cypermethrin, deltamethrin, lambda-cyhalothrin) as reported by Gunning *et al*., 1984; Armes *et al*., 1996; McCaffery, 1998. Over-dependence on a particular group of chemicals is one of the important reasons for the rapid development of resistance. With this development of insecticide resistance, the control of *H. armigera* has become critical in many regions worldwide (Tabashnik*et al*., 2014). The resistance vary based on insecticide usage patterns which is dependent on host crop and growing conditions (Kranthi*et al*.,2001; IndraChaturvedi, 2007; Sagar*et al*., 2013;Dilbar*et al*., 2014; Honnakerappaand Udikeri, 2018).In Karnataka, India different host crops of *H*.*amigera* are largely under different agroclimaticconditions representing different soils, precipitation pattern and irrigation facility as well. Thus quite ideal situations offering varied degree of resistance in any pest are available in Karnataka (Table 1).So, to know information about the last visibility of insecticide resistance to *H. armigera*due to changes in pesticide usage pattern, after introduction of Bt cotton and influential pest outbreak, in addition to that limited insight of resistance on major cropping systems for Helicoverpa in different agro-ecological zones of Karnataka.

**Table 1:** Rainfall, crop dominance, type of soil and usage of insecticide in agro climatic zones of Karnataka, India.

| **SL. No** | **Agro Climatic Zones** | **Annual Rainfall (mm)** | **Soil Type** | **Principal Crops** | **Insecticide Usage Level** |
| --- | --- | --- | --- | --- | --- |
| 1 | North East Transition Zone | 830-890 | Major - clay,<br>Minor -Laterite | Pulses, Jowar, Oilseeds, Bajra, Cotton and Sugarcane | High |
| 2 | North-eastern Dry Zone | 633.2 to 806.6 | Major - deep black clay<br>Minor - medium black | Rabi jowar, Bajra, Pulses, Oilseed | High |
| 3 | Northern Dry Zone | 464.5-785.7 | Major - shallow to deep black clay | Rabi jowar, Maize, Bajra, Groundnut, Cotton, Wheat, Sugarcane and Tobacco | Moderate |
| 4 | Central Dry Zone | 453.5-717.7 | Major - red sandy loams<br>Minor - shallow to deep black | Ragi, Jowar, Pulses and Oilseeds | Low |
| 5 | Eastern Dry Zone | 679.1-888.9 | Major - red loamy<br>Minor - lateritic | Ragi, Rice, Pulses, Maize and Oil seeds | Moderate |
| 6 | Southern Dry Zone | 670.6-888.6 | Major - red sandy loam<br>Minor - red loamy | Rice, Ragi, Pulses, Jowar and Tobacco | Moderate |
| 7 | Southern Transition Zone | 611.7-1053.9 | Major - red sandy loam<br>Minor - red loamy | Rice, Ragi, Pulses, Jowar and Tobacco | Low |
| 8 | North Transition Zone | 619.4-1303.2 | Black clay and red sandy loam | Rice, Jowar, Groundnut, Pulses, Sugarcane and Tobacco | Low to Moderate |
| 9 | Hill Zone | 904.4-3695.1 | -- | rice and pulses | Low |
| 10 | Coastal Zone | 3010.9-4694.4 | Red lateritic and coastal alluvial<br>Black clay and red sandy loam | Rice, Pulses and Sugarcane | Low |

## 2. Materials and Methods

The experiment was conducted at the Agriculture Research Station, Dharwad (Hebballi) farm of University of Agricultural Sciences, Dharwad during 2017 in laboratory to know the situation of the resistant scenario of *H. armigera* in different agroclinatic zones of Karnataka. The *H. armigera* larvae were collected from different locations of Karnataka (Fig. 1), representing different cropping patterns and agro-ecosystems.

**Fig. 1.**
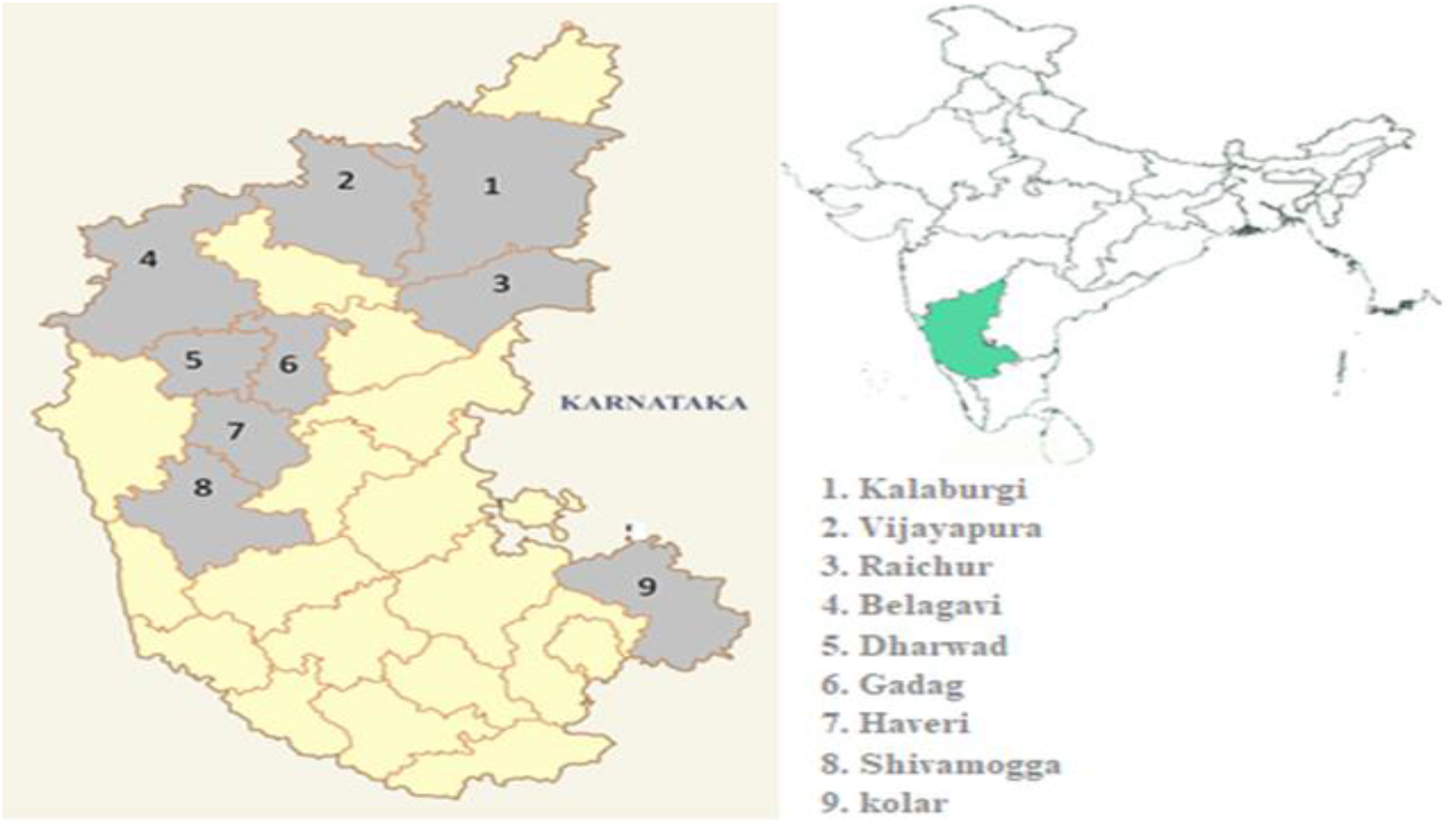
Locations selected for collection of *H. armigera* for resistance study in Karnataka.

### 2.1 Collection of *H. armigera* populations

The *H. armigera* larvae were collected from different locations of the Karnataka (Fig. 1) representing different cropping patterns and agro-ecosystems during 2017, i.eRaichur and Kalburgi come under the Northeastern dry zone, Vijayapur and Gadag come under the Northern dry zone, Dharwad, Haveri and Belagavi come under North transition zone, Shivamogga Southern transition zone and Kolar comes under Eastern dry zone in such way that it showing diverse population belong different climatic conditions and cropping patterns of Karnataka (Table 3). These populations were assigned with the strain codes according to locations. Then each strain was reared on an artificial diet for the next generations (F1)for bioassys

### 2.2 Artificial diet preparation for *H. armigera*

The artificial diet was prepared as per the procedure outlined by Kranthi (2005). The rearing was as per methods prescribed by Ahmad *et al*. (2003) Neonate larvae were transferred to plastic containers containing an artificial diet. Then before reaching the second instar larvae were transferred to multiwell rearing (25 or 24 wells) trays and maintained until pupation. The larvae were shifted to trays having a fresh diet every third day.

### 2.3 Maintenance of pupae and adult

Pupae obtained from each field-collected larva were treated with surface sterile solution 0.5-1 % of sodium hypochlorite, gently washed with distilled water and moisture was removed with help of tissue paper and air-dried for a few minutes. Later pupae were kept for adult emergence in a wooden cage 9.2” × 9.2” × 9.3” having mesh on three sides and glass in front. The pupae were kept in bread boxes having moist sand and sawdust (1:1) mixture.

The emerged adults were allowed to feed on an adult diet and one or two pairs of adults (male and female) were kept in an egg cage for mating and egg-laying. For an adult diet, 5 gm each of sucrose and honey was dissolved in 90 ml of sterile water and boiled for 5 minutes. After proper cooling, 0.2 g each of ascorbic acid and methyl hydroxypara benzoate were added and stored at 4.0 ^0^C for 1-2 weeks (Kranthi, 2005). Sterile absorbent cotton swabs were soaked in the adult diet solution and placed in a cage for adult feeding which was changed on alternate days. The cage was covered with a fine black muslin cloth. The eggs laid on muslin cloth were removed with a camel hair brush and dipped in 0.1 % sodium hypochloride surface sterile solution. These eggs were placed in small plastic jars for hatching. A temperature of 27 ± 1 °C with a photoperiod of 14D: 10L and relative humidity of 65 ± 5% was maintained.

### 2.4 F_1_ population maintenance

The larvae that emerged in each population were maintained in groups (first instar only) on an artificial diet in large size plastic containers having mesh lids. Further larvae were maintained individually in 25 or 24 well trays having an artificial diet till reaching third instars. The third instar larvae were subjected to bio-assays.

### 2.5 Susceptible strain

An insecticide susceptible laboratory strain of *H. armigera*furnished by Dr.VinayKalia, Professor, Division of Entomology, IARI New Delhi (as gratis) was maintained separately in an artificial diet as explained earlier.

### 2.6 Determination of insecticide resistance in *H. armigera*

#### 2.6.1 Test insecticides

The insecticides used in the study and formulations have been presented in Table 2. All insecticides were obtained as commercial formulations available in the market during 2016. The required concentrations of test insecticides were prepared from the formulated products by dissolving the required quantities in double distilled water after accurate weighment. The solution thus prepared was preserved in the refrigerator for further use. Concentrations limiting the mortality between 10-90% were used for probit, log dose mortality analysis. To arrive at five test concentrations, a couple of round pilot tests were carried out.

**Table 2.**
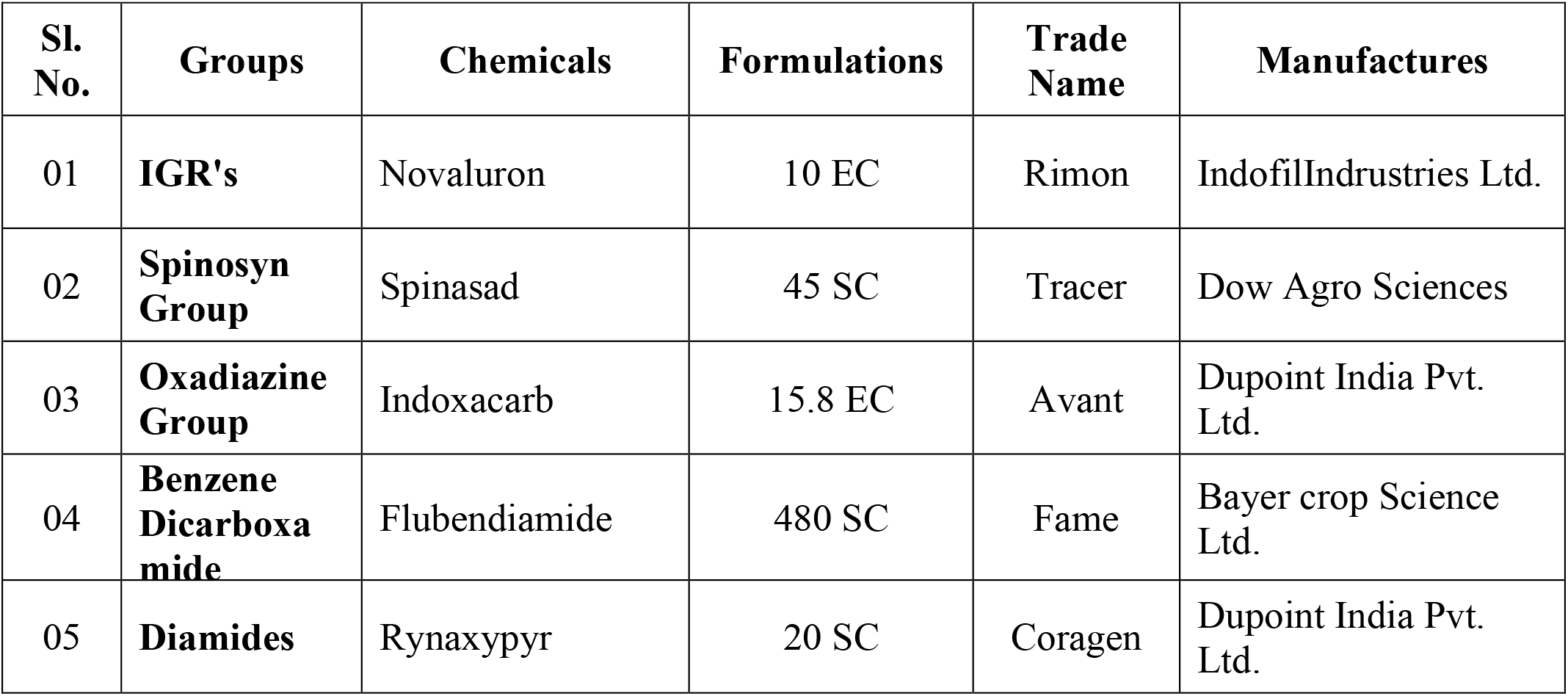
Insecticides used for the determination of resistance in *H. armigera*.

**Table 3.**
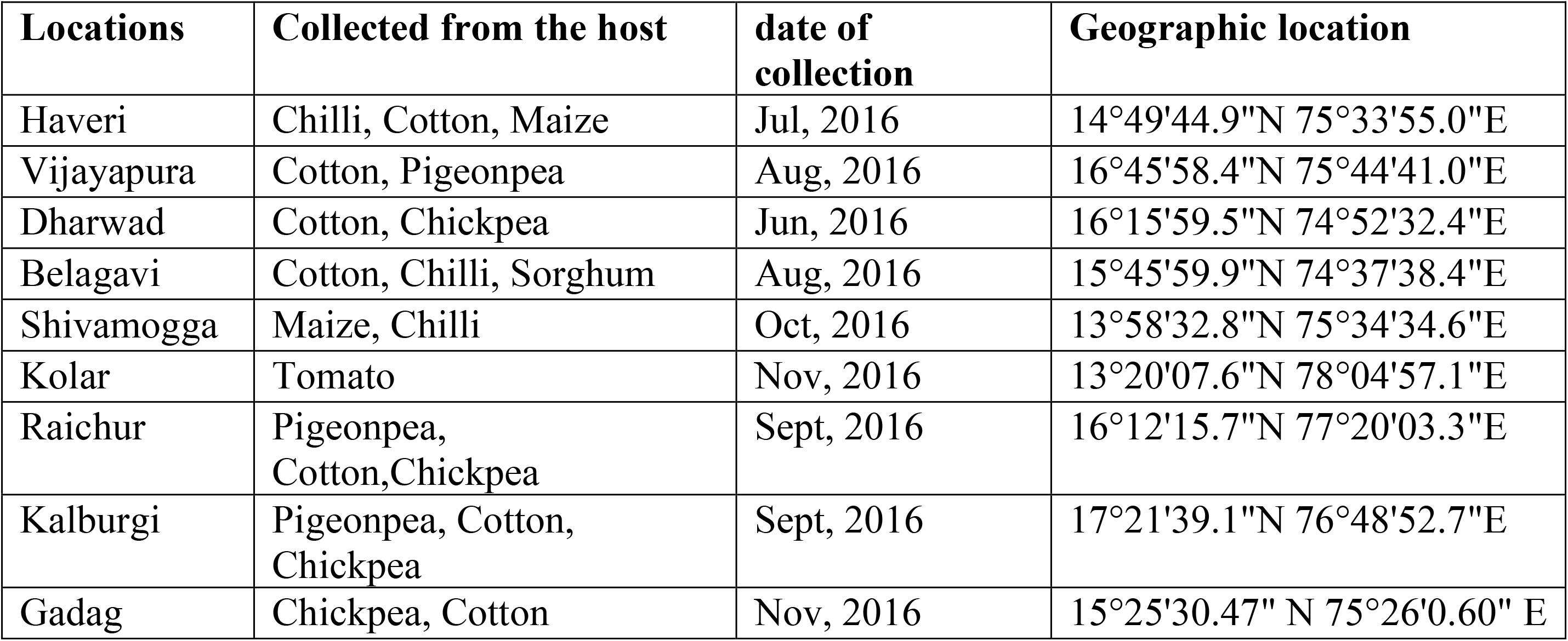
*H. armigera* collected from different location for resistance study.

#### 2.6.2 Bioassays

The resistance in different field strains of *H. armigera* and the susceptible strain was assessed through the leaf disc method as suggested for specific insecticides (Singh and Mahal, 2005). Fully expanded DCH-32 Non-Bt cotton leaves grown in insecticide-free conditions separately were used for this purpose. From such leaves, 5.0 cm diameter discs were cut open from the central part of such leaves using a metal borer. Then these discs were dipped into different concentrations of the insecticides for 10 seconds and dried for 30 minutes (Rafiee*et al*., 2008) in air. These leaf discs were placed into plastic Petri dishes lined with moistened filter paper to avoid desiccation.Ten third instar larvae from F1 generation larvae with an average weight of 30 ± 3.0 mg were used for every concentration of each insecticide. Assay concentration was replicated for 4 times. A treatment with leaf disc immersed in distilled water served as control.

#### 2.6.3 Data collection

Larval mortality was recorded at 24, 48 and 72 hours after treatment. The mortality percent at 72 hours after treatment was considered as the endpoint for the assessment of toxicity of test insecticides (Fisk and Wright, 1992). The mortality was determined based on the failure of insects to move upon coordinated pronding.

#### 2.6.4 Data analysis and interpretation of resistance levels

The larval mortality percent at 72 hours after treatment was considered as the endpoint for the assessment of toxicity of test insecticides (Fisk and Wright, 1992). Results were expressed as percentage mortality using Abbott’s formula. Data were analyzed by probit analysis (Finney, 1972) with SPSS statistical computer programme. To determine cross-resistance among the insecticides tested, pair wise correlation coefficients of log LC50 values of the common populations for each insecticide were calculated by SPSS statistical computer program and for better graphical Interpretation R studio software was used.

The mortality data was subjected for correction using Abbott’s formula

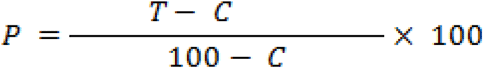

Where, P = The corrected mortality percentage

T = The observed percentage mortality in the treatment

C = Thepercent of mortality in the control

Then LC_50_ wasdeveloped by probit analysis (Finney, 1972) method. Further resistance was assessed by the working resistance ratio for each insecticide and strain.

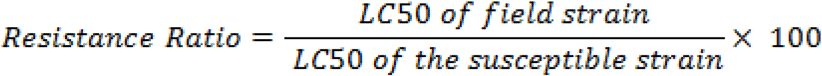

### 2.7 Morphometric parameters of *H. armigera*strains

Morphometric variations of larvae, pupae and adults of different strains were recorded inF1 population. To study the larval morphometric characters length and weight of fifty larvae per strain where considered. General body colours of the larval stage were recorded by visual observation. The length of the fifth instar larva was measured by using a centimeter-scale while microbalance was used for recording weight. Pupal length and weight were recorded for 20 randomly selected pupae of *H. armigera* from each location. Pupal length (mm) and weight (mg) were measured by centimeter-scale and microbalance, respectively (Fakruddin*et al*, 2007).

### 2.8 Collection of information on insecticide usage pattern and selection pressure against *H. armigera*

Insecticide usage patterns in each region or cropping system were unearthed through interviews with farmers using a structured schedule. The extent of exposure to different types of pesticides dosages, frequency and other pest management practices targeting *H. armigera* (including refugia in Bt cotton) was addressed. In each district representing a particular region, 50 farmers were selected for an interview to know pest management practices prevailing amongst them. Thus a total of 450 farmers were consulted during the survey. The data collection was on a questionnaire yes or no based on particular aspects which could help draw a conclusion about resistance over tested insecticides in different locations of Karnataka. The data gathered was catered to MS-Excel master worksheet, sorted and manifested.

To interpret cross-resistance spectra among the insecticides tested, pairwise correlation coefficients of LC50 values of the common populations for each insecticide (Ahmad *et al*., 2006) were calculated by R software.

## 3. Results

### 3.1 IGR and newer insecticides resistance in field populations of *H. armigera*

#### 3.1.1 IGRs (Nuvaluron) resistance in

The RCR strain recorded the highest resistance with LC50 value of 18.07 ppm followed by KBG (16.25 ppm) and HVR (13.42 ppm) strains. The lowest resistance was observed in KLR (10.86 ppm). Hence resistance ratio was maximal for RCR (1.95 fold), KBG (1.75 fold) and HVR (1.45 fold) strains and the least resistance ratio was for KLR (1.17 fold) strain in Nuvaluron (Table 4).

**Table 4.** Toxicity ofIGR, *Btk*and newer insecticides against different strains of *H. armigera* fromvarious localities in Karnataka during 2016-17.

| CHEMICALS | Strain | LC50 (PPM) | FL 95% | LC90 (PPM) | SLOPE ± SE | RR |
| --- | --- | --- | --- | --- | --- | --- |
| **Novaluron** | SUS-L | 9.27 | 6.4-12.02 | 48.26 | 1.78±0.28 | - |
| | HVR | 13.42 | 9.05-18.31 | 114.98 | 1.37±0.23 | 1.45 |
| | VJP | 13.02 | 8.65-17.87 | 115.89 | 1.35±0.23 | 1.4 |
| | DWD | 13.25 | 8.74-18.30 | 124.34 | 1.31±0.23 | 1.43 |
| | BLG | 12.85 | 8.33-17.84 | 125.52 | 1.29±0.23 | 1.39 |
| | SMG | 11.75 | 7.82-15.95 | 92.77 | 1.42±0.24 | 1.27 |
| | KLR | 10.86 | 7.05-14.84 | 87.3 | 1.41±0.23 | 1.17 |
| | RCR | 18.07 | 13.28-24.2 | 127.57 | 1.51±0.24 | 1.95 |
| | KBG | 16.25 | 11.95-21.44 | 105.68 | 1.57±0.24 | 1.75 |
| | GDG | 12.62 | 8.25-17.42 | 116.8 | 1.32±0.23 | 1.36 |
| **Spinasad** | SUS-L | 9.03 | 6.51-11.46 | 42.41 | 1.9±0.30 | - |
| | HVR | 9.59 | 6.44-12.73 | 64.28 | 1.55±0.27 | 1.06 |
| | VJP | 9.39 | 2.39-16.56 | 52.02 | 1.72±0.27 | 1.04 |
| | DWD | 9.78 | 6.58-12.98 | 66.35 | 1.54±0.27 | 1.08 |
| | BLG | 9.56 | 6.7-12.41 | 53.58 | 1.71±0.27 | 1.06 |
| | SMG | 9.67 | 6.72-12.62 | 56.73 | 1.66±0.27 | 1.07 |
| | KLR | 9.85 | 6.86-12.86 | 58.45 | 1.65±0.27 | 1.09 |
| | RCR | 9.9 | 6.6-13.22 | 70.76 | 1.50±0.26 | 1.1 |
| | KBG | 10.19 | 6.91-13.53 | 70.45 | 1.52±0.26 | 1.13 |
| | GDG | 9.15 | 6.36-11.91 | 50.95 | 1.71±0.27 | 1.01 |
| **Indoxacarb** | SUS-L | 5.64 | 3.41-8.11 | 57.06 | 1.27±0.19 | - |
| | HVR | 6.14 | 3.7-8.90 | 69.76 | 1.18±0.19 | 1.12 |
| | VJP | 6.31 | 3.88-9.01 | 65.48 | 1.26±0.19 | 1.15 |
| | DWD | 6.58 | 3.95-9.60 | 81.93 | 1.17±0.19 | 1.20 |
| | BLG | 6.63 | 4.02-9.62 | 79.51 | 1.18±0.19 | 1.21 |
| | SMG | 5.94 | 3.53-8.64 | 68.75 | 1.2±0.19 | 1.08 |
| | KLR | 6.09 | 3.63-8.87 | 71.76 | 1.19±0.19 | 1.11 |
| | RCR | 6.96 | 4.07-10.35 | 103.83 | 1.09±0.18 | 1.27 |
| | KBG | 6.70 | 3.99-9.83 | 88.59 | 1.14±0.19 | 1.22 |
| | GDG | 6.51 | 3.99-9.4 | 73.77 | 1.21±0.19 | 1.18 |
| **Flubendiamide** | SUS-L | 3.57 | 2.60-4.47 | 15.17 | 2.04±0.33 | - |
| | HVR | 3.77 | 2.82-4.65 | 14.92 | 2.14±0.33 | 1.06 |
| | VJP | 3.64 | 2.60-4.59 | 16.66 | 1.93±0.32 | 1.02 |
| | DWD | 3.74 | 2.73-4.68 | 16.44 | 1.99±0.33 | 1.05 |
|  | BLG | 3.72 | 2.69-4.67 | 16.74 | 1.96±0.30 | 1.04 |
|  | SMG | 3.63 | 2.65-4.54 | 15.51 | 2.03±0.33 | 1.02 |
|  | KLR | 3.66 | 2.64-4.6 | 16.37 | 1.97±0.33 | 1.03 |
|  | RCR | 3.82 | 2.82-4.77 | 16.51 | 2.01±0.33 | 1.07 |
|  | KBG | 3.66 | 2.64-4.6 | 16.37 | 1.97±0.33 | 1.03 |
|  | GDG | 3.65 | 2.68-4.55 | 15.25 | 2.06±0.33 | 1.02 |
| **Rynaxypyr** | SUS-L | 4.95 | 2.97-7.11 | 51.19 | 1.26±0.20 | - |
|  | HVR | 5.26 | 3.12-7.64 | 61.12 | 1.2±0.20 | 1.06 |
|  | VJP | 5.13 | 3.13-7.32 | 51.69 | 1.27±0.20 | 1.04 |
|  | DWD | 5.31 | 3.28-7.53 | 52.18 | 1.29±0.20 | 1.07 |
|  | BLG | 5.23 | 3.17-7.5 | 54.97 | 1.25±0.20 | 1.06 |
|  | SMG | 5.19 | 3.12-7.47 | 56.15 | 1.24±0.20 | 1.05 |
|  | KLR | 5.01 | 2.96-7.26 | 55.66 | 1.22±0.22 | 1.01 |
|  | RCR | 5.38 | 3.29-7.69 | 56.62 | 1.25±0.20 | 1.09 |
|  | KBG | 5.34 | 3.33-7.55 | 51.14 | 1.3±0.20 | 1.08 |
|  | GDG | 5.16 | 3.18-7.34 | 50.65 | 1.29±0.20 | 1.04 |

#### 3.1.2 Newer insecticides

The maximal resistance level was found with a lethal concentration in RCR strain 6.96 ppm and followed by KBG strains 6.7 ppm in Indoxacarb with (1.27 fold and 1.22 fold) respectively, whereas, in Rynaxypyr 5.38 ppm in RCR strain, and 5.34 ppm in KBG strain and the resistance ratio 1.09 fold and 1.08 fold respectively, similarly for Spinasad is KBG strain 10.19 ppm and followed by RCR9.90 ppm lethal concentration and resistance ratio (1.13 fold and, 1.10 fold), respectively in the Flubendiamide maximal lethal concentration in RCR strain 3.82 ppm, 1.07 fold resistance ratio followed by HVR strain 3.77 ppm lethal concentration and resistance ratio 1.07 fold.

The merest resistance level was recorded in GDG strain 9.15 ppm and resistance ratio 1.01 fold for Spinasad, whereas, in Indoxacarb the KLR strain 6.09 ppm and resistance ratio 1.11 fold and 5.01 ppm lethal concentration and resistance ratio 1.08 fold seen in KLR strain for Rynaxypyr, similarly in case ofFlubendiamide significant subordinate resistance were weird in HVR, SMG and GDG strains (Table 4).

Paired comparisons of the log LC50s for the same populations for the different insecticides (Table 5) positive correlation was perceived in all the insecticides which have been tested and significant positive correlation (0.71) subsist only between Indoxacarb and Rynaxypyr and positive correlation found in Novaluron, Spinasad and Flubendiamide.

**Table 5.** Pairwise correlation coefficient comparisons between log LC50 Insecticides tested on the field populations of *Helicoverpaarmigera*.

|  | Novaluron | Spinasad | Indoxacarb | Flubendiamide | Rynaxypyr |
| --- | --- | --- | --- | --- | --- |
| Novaluron | 1 |  |  |  |  |
| Spinasad | 0.488 | 1 |  |  |  |
| Indoxacarb | 0.805 | 0.266 | 1 |  |  |
| Flubendiamide | 0.598 | 0.242 | 0.533 | 1 |  |
| Rynaxypyr | 0.847 | 0.44 | .701* | 0.648 | 1 |

\*. Correlation is significant at the 0.05 level (2-tailed).

#### 3.1.3 Morphometric parameters of *H. armigera* across geographical populations of Karnataka along with susceptible lab strain

In susceptible lab strain larval length in range of 1.85-2.50 cm with mean 2.48±0.41 cm and larval weight in the range of 0.395-0.500 g with mea 0.423±0.15 gm, correspondingly pupal length and weight in range of 1.30-1.85 cm and 0.17-0.30 gm with mean 1.58±0.14 cm and 0.225±0.03 g respectively.

RCH strain from Raichur location was preeminent in all the morphometric parameters range and mean of larval length (2.00-3.25 cm and 2.75±0.48 cm), larval weight (0.460-0.600 g and 0.511±0.04 g),pupal length (1.50-2.00 cm and 1.76±0.18 cm) and pupal weight (0.22-0.39 g and 0.309±0.05 g) which was chased by KLB strain from Kalaburgi with range and mean larval length (2.00-3.20 cm and 2.71±0.42 cm), larval weight (0.470-0.590 g and 0.462±0.17 g), pupal length (1.50-2.00 cm and 1.72±0.20 cm) and pupal weight (0.23-0.38 g and 0.286±0.05 g). Apart from the susceptible strain the minimal morphometric parameters were made out in SMG strain from Shivamogga as for range and mean larval length (1.90-2.95 cm and 2.52±0.49 cm) larval weight (0.400-0.520 g and 0.454±0.04 g), pupal length (1.35-1.90 cm and 1.62±0.16 cm) and pupal weight (0.18-0.30g and 0.237±0.03 g) which was trailed by KLR strain from Kolar location with a range of larval length and weight (1.90-2.95 cm and 0.400-0.550 g) and mean larval length and weight (2.58±0.47 cm and 0.462±0.06 g) and also a range of pupal length and weight (1.35-1.90 cm and 0.201-0.352 g) and mean of pupal length and weight (1.63±0.15 cm and 0.239±0.03 g). The remaining strains do not menifesting much disparity which fits into the above mentioned categories of superior and inferior (Table 6).

**Table 6.**
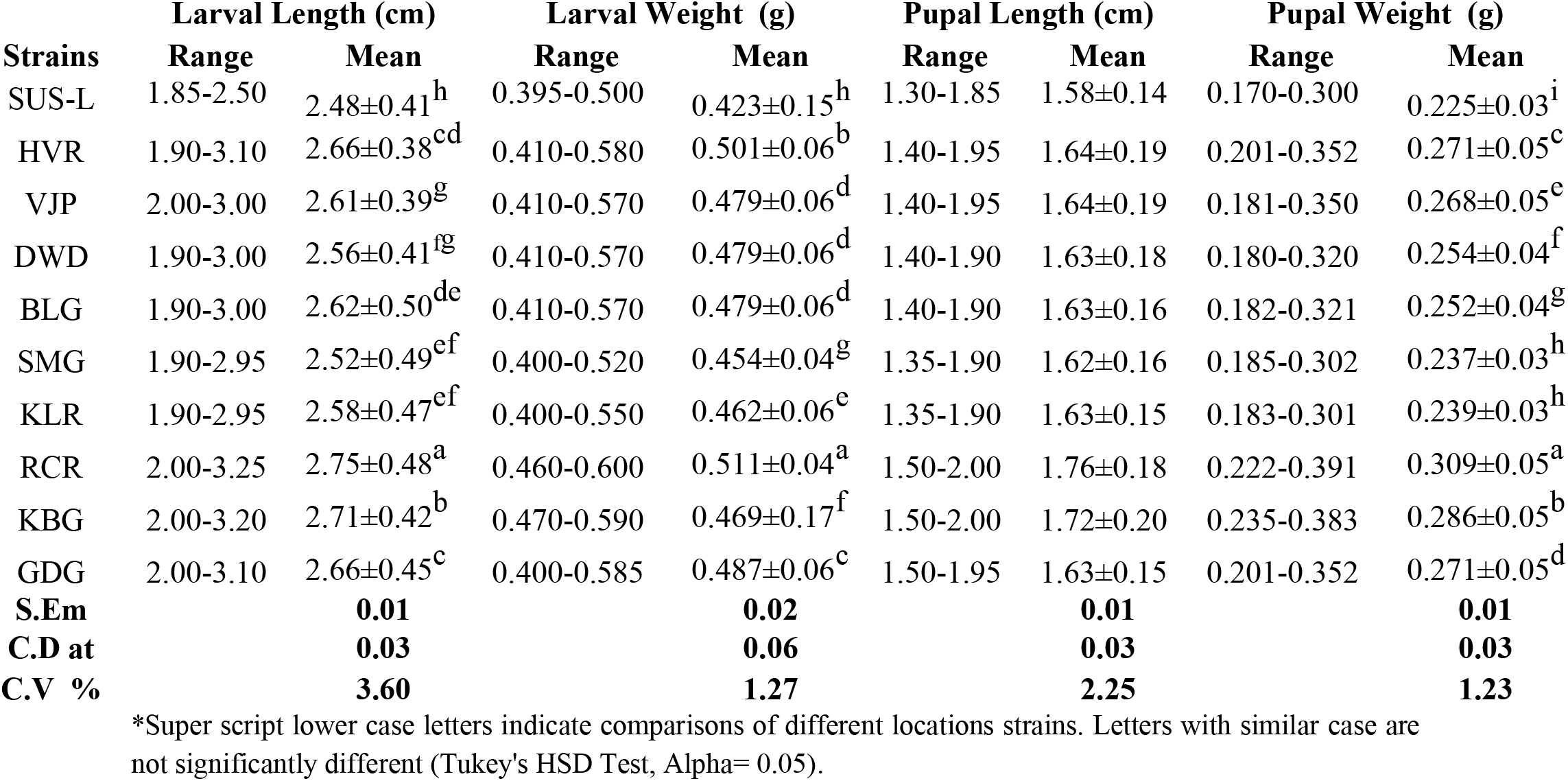
Length and Weigth of larva and its pupa of *H. armigera* across geographical populations of Karnataka.

| Strains | Larval Length (cm) |  | Larval Weight (g) |  | Pupal Length (cm) |  | Pupal Weight (g) |  |
| --- | --- | --- | --- | --- | --- | --- | --- | --- |
|  | Range | Mean | Range | Mean | Range | Mean | Range | Mean |
| SUS-L | 1.85-2.50 | 2.48±0.41<sup>h</sup> | 0.395-0.500 | 0.423±0.15<sup>h</sup> | 1.30-1.85 | 1.58±0.14 | 0.170-0.300 | 0.225±0.03<sup>i</sup> |
| HVR | 1.90-3.10 | 2.66±0.38<sup>cd</sup> | 0.410-0.580 | 0.501±0.06<sup>b</sup> | 1.40-1.95 | 1.64±0.19 | 0.201-0.352 | 0.271±0.05<sup>c</sup> |
| VJP | 2.00-3.00 | 2.61±0.39<sup>g</sup> | 0.410-0.570 | 0.479±0.06<sup>d</sup> | 1.40-1.95 | 1.64±0.19 | 0.181-0.350 | 0.268±0.05<sup>e</sup> |
| DWD | 1.90-3.00 | 2.56±0.41<sup>fg</sup> | 0.410-0.570 | 0.479±0.06<sup>d</sup> | 1.40-1.90 | 1.63±0.18 | 0.180-0.320 | 0.254±0.04<sup>f</sup> |
| BLG | 1.90-3.00 | 2.62±0.50<sup>de</sup> | 0.410-0.570 | 0.479±0.06<sup>d</sup> | 1.40-1.90 | 1.63±0.16 | 0.182-0.321 | 0.252±0.04<sup>g</sup> |
| SMG | 1.90-2.95 | 2.52±0.49<sup>ef</sup> | 0.400-0.520 | 0.454±0.04<sup>g</sup> | 1.35-1.90 | 1.62±0.16 | 0.185-0.302 | 0.237±0.03<sup>h</sup> |
| KLR | 1.90-2.95 | 2.58±0.47<sup>ef</sup> | 0.400-0.550 | 0.462±0.06<sup>e</sup> | 1.35-1.90 | 1.63±0.15 | 0.183-0.301 | 0.239±0.03<sup>h</sup> |
| RCR | 2.00-3.25 | 2.75±0.48<sup>a</sup> | 0.460-0.600 | 0.511±0.04<sup>a</sup> | 1.50-2.00 | 1.76±0.18 | 0.222-0.391 | 0.309±0.05<sup>a</sup> |
| KBG | 2.00-3.20 | 2.71±0.42<sup>b</sup> | 0.470-0.590 | 0.469±0.17<sup>f</sup> | 1.50-2.00 | 1.72±0.20 | 0.235-0.383 | 0.286±0.05<sup>b</sup> |
| GDG | 2.00-3.10 | 2.66±0.45<sup>c</sup> | 0.400-0.585 | 0.487±0.06<sup>c</sup> | 1.50-1.95 | 1.63±0.15 | 0.201-0.352 | 0.271±0.05<sup>d</sup> |
| **S.Em** |  | **0.01** |  | **0.02** |  | **0.01** |  | **0.01** |
| **C.D at** |  | **0.03** |  | **0.06** |  | **0.03** |  | **0.03** |
| **C.V %** |  | **3.60** |  | **1.27** |  | **2.25** |  | **1.23** |

\*Super script lower case letters indicate comparisons of different locations strains. Letters with similar case are not significantly different (Tukey's HSD Test, Alpha= 0.05).

A correlation exists between morphometric parameters two groups of insecticide which were tested in the study *viz*, IGR group Novaluron and Newer insecticides in Table 7, positive correlation discerns in both groups of insecticides with the larval weight and pupal length as well as weight, alike significant positive correlation (0.781* and 0.681*) respectively, be the case with both insecticidal group with a larval length of *H. armigera*.

**Table 7.**
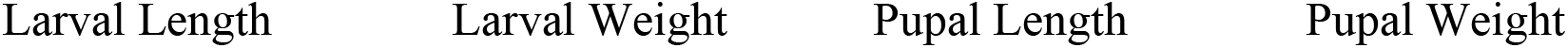
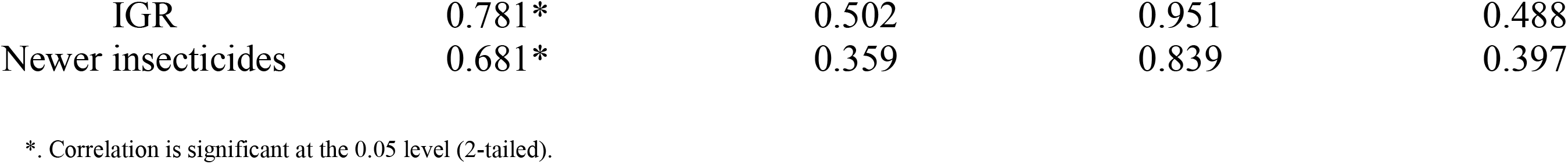
Pairwise Correlation between morphometric parameters and Insecticides resistance levels on the field populations of *Helicoverpaarmigera*.

#### 3.1.4 Pesticide usage pattern and selection pressure about *H. armigera* in different locations of Karnataka

According to a survey conducted during 2016-17 farmers from different locations of Karnataka (Table 8), 31.67 percent were aware of recommended pesticides against *H. armigera*if we consider social status 62.50 percent of illiterate and 41.67 percent of educated farmers were seen in different locations pertaining to the growers of the major crop which were selected for experiment for managing *H. armigera*.

**Table 8.**
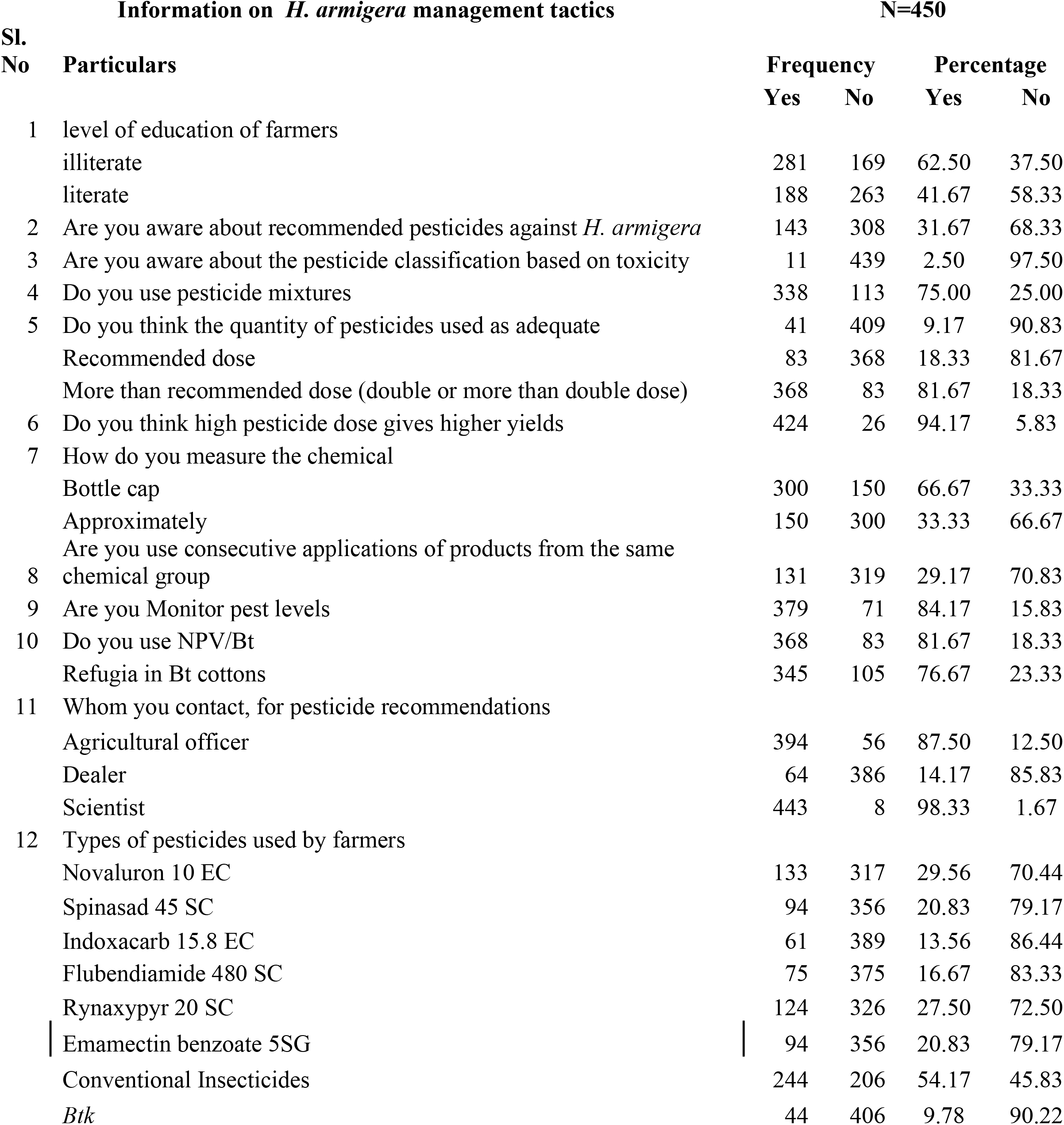
Usage pattern of insecticides to manage *H*.*armigera*in Karnataka,India.

2.5 percent of farmers were appreciative pertaining to the pesticide classification based on toxicity but 75 percent of farmers were well versed with pesticide mixture, whereas the adequate quantity of pesticide usage according to recommended dosage was 18.33 and more than recommended dose was 81.67 percent because 95.83 percent of farmers were thinking that high pesticide dose would give higher yield, however for pesticide recommendations farmers used to contact more of dealers (85.83 percent) and less of agriculture officers (12.50 percent) and scientist (1.67 percent). Most of the farmers about 66.67 percent were measuring the chemical approximately and 33.33 percent were by bottle cap that may be due to the level of education 62.50 percent Famers come under illiterate category and 37.50 percent literate which varied from primary school to graduation.

A most important aspect of resistance management to avoiding consecutive applications of insecticides from the same chemical group only 29.17 percent are avoided, hardly 15.83 percent, 18.33 percent and 23.33 percent of farmers were practicing monitoring pest level through sex pheromone traps, use of NPV/Bt and refugial strategies, respectively.

The proportion of different groups of insecticides used by the farmers against *H. armigera* (Table 8) where 54.17 percent of conventional insecticides and newer insecticides which were tested in this experiment were used by farmers Spinasad 20.83 percent, Indoxacarb 13.56 percent, Flubendiamide 16.67 percent, Rynaxypyr 27.50 percent, Emamectin benzoate 20.83 percent, 29.56 percent IGR (Novaluron) and 9.78 percent*Btk*.

## 4. Discussion

Though newer insecticides having different modes of action have a significant role in the pest management arena, the strains *H. armigera* have shown varying responses. Among nine locations the maximum resistance was found to be in RCR strain against newer chemistry insecticides *viz*., Indoxacarb (1.27 fold), Flubendiamide (1.07 fold) and Rynaxypyr (1.09 fold), alike KBG strain showed maximum Spinosad (1.13 fold) resistance. Minimal resistance was noted in SMG, VJP and GDG strains. Newer chemicals have got less resistance as compared with the Novaluran in different strains of Karnataka as shown in Fig. 2. Maximum resistance was noticed in RCR (1.95 fold) followed by KBG (1.75 fold) and HVR strain (1.45 fold) against a least in KLR (1.17 fold) and other strains *viz*., SMG and GDG against Nuvaluron.

**Fig. 2.**
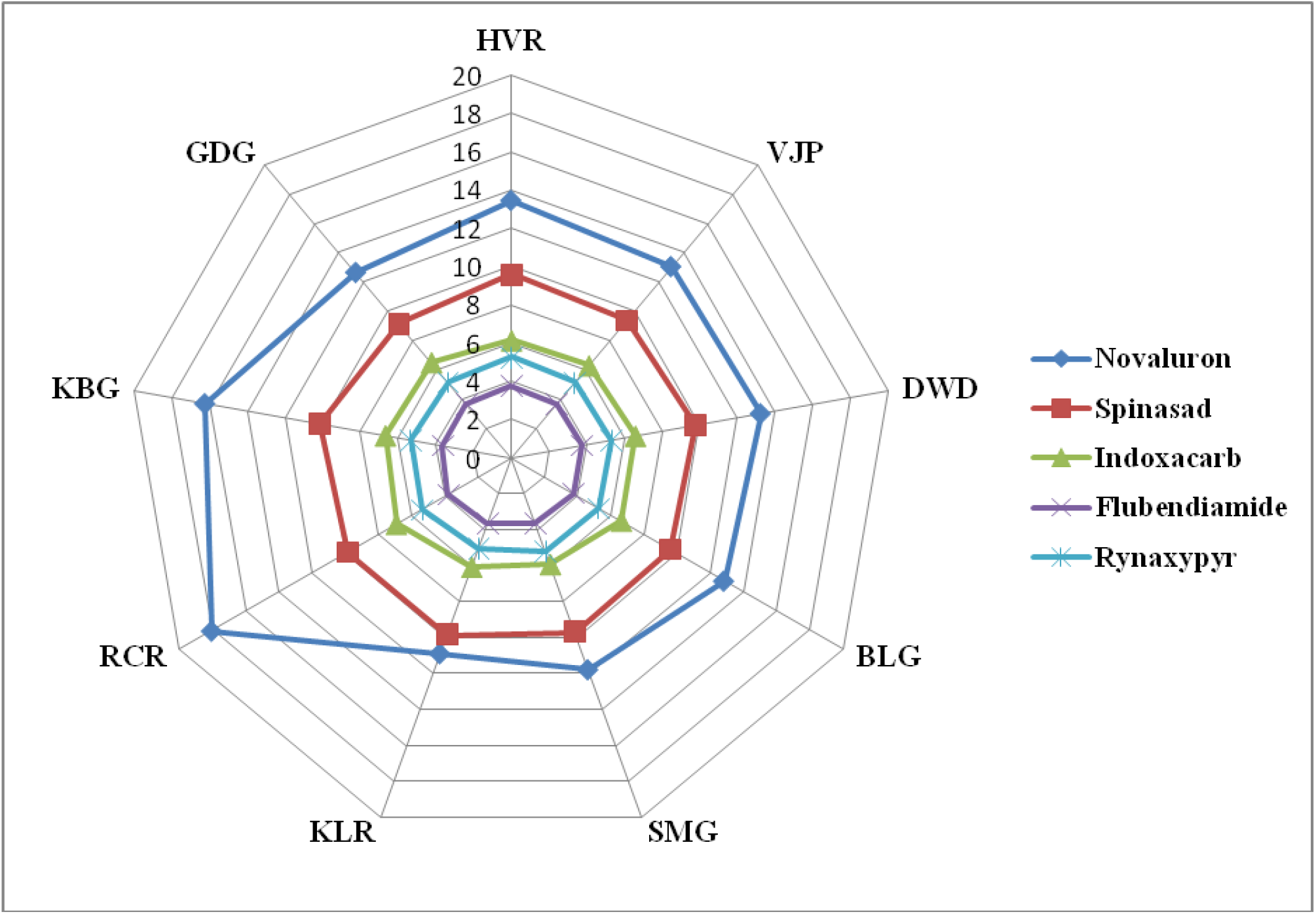
Resistance level of selected insecticides against field populations of Karnataka.

The present study indicates a negligible resistance and variability in *H. armigera* strains to newer molecules. Similarly, Mirza Abdul *et al*. (2015) observed lower lethal concentrations of these new chemicals compared to OP’s and pyrethroids having higher LC_50_ values to *H. armigera*. Further Kranthi*et al*. (2000), Rao (2008), Cook*et al*. (2005) and Gupta *et al*. (2005) have registered a negligible variation sensitively, but least lethal concentration requirement against *H. armigera* with respect to spinosad and indoxacarb. Mushtaq and Rashid (2015) also have reported the very low-level resistance to growth regulators owing to their novel modes of action. In India, IGR resistance is rarely addressed.

The use of such newer chemicals would check the pyrethroids resistance as reported by SufianSaif*et al*., 2013. Thus these insecticides could be considered as most effective and safe in *H. armigera*management throughout Karnataka. These results of LC_50_ represent initial efforts to develop baseline data for Rynoxypyr, Flubendiamide, Spinosad and Indoxacarb as so far no such studies have been done considering these populations although the use of leaf disc method bioassays may not provide the optimum scale of the toxicity for all the tested insecticticides, the procedure appeared to perform well for those insecticides that require ingestion. This baseline data should assist in monitoring for changes in susceptibility to these new insecticides as their use becomes widespread across multiple host crops in Karnataka.

From correlation analysis among different insecticides clearly depicts the strong relation of Novaluron with the newer insecticides which have been tested; a significant positive correlation exists between Rynaxypyr and Indaxocarb. However, newer insecticides interestingly Spinosad showed compact correlation with other tested insecticides if observed in Fig. 3 network corrplot showing separate network pathway and less intensity of color away from other insecticides, in this finding among newer insecticides may Spinosad undoubtedly hope for rotation in governing insecticide resistance against *H. armigera*even though newer insecticides are a better choice for resistance management. Rotation of still-effective conventional chemistries with the new chemistries such as emamectin benzoate and insect growth regulators (IGRs), novaluron, is recommended (Ahmad *et al*., 2006).So, comprehensively stipulates that we should steer clear of using different groups of insecticides for resistance management in order to avoid the menace of *H. armigera*in different cropping systems.

**Fig. 3.**
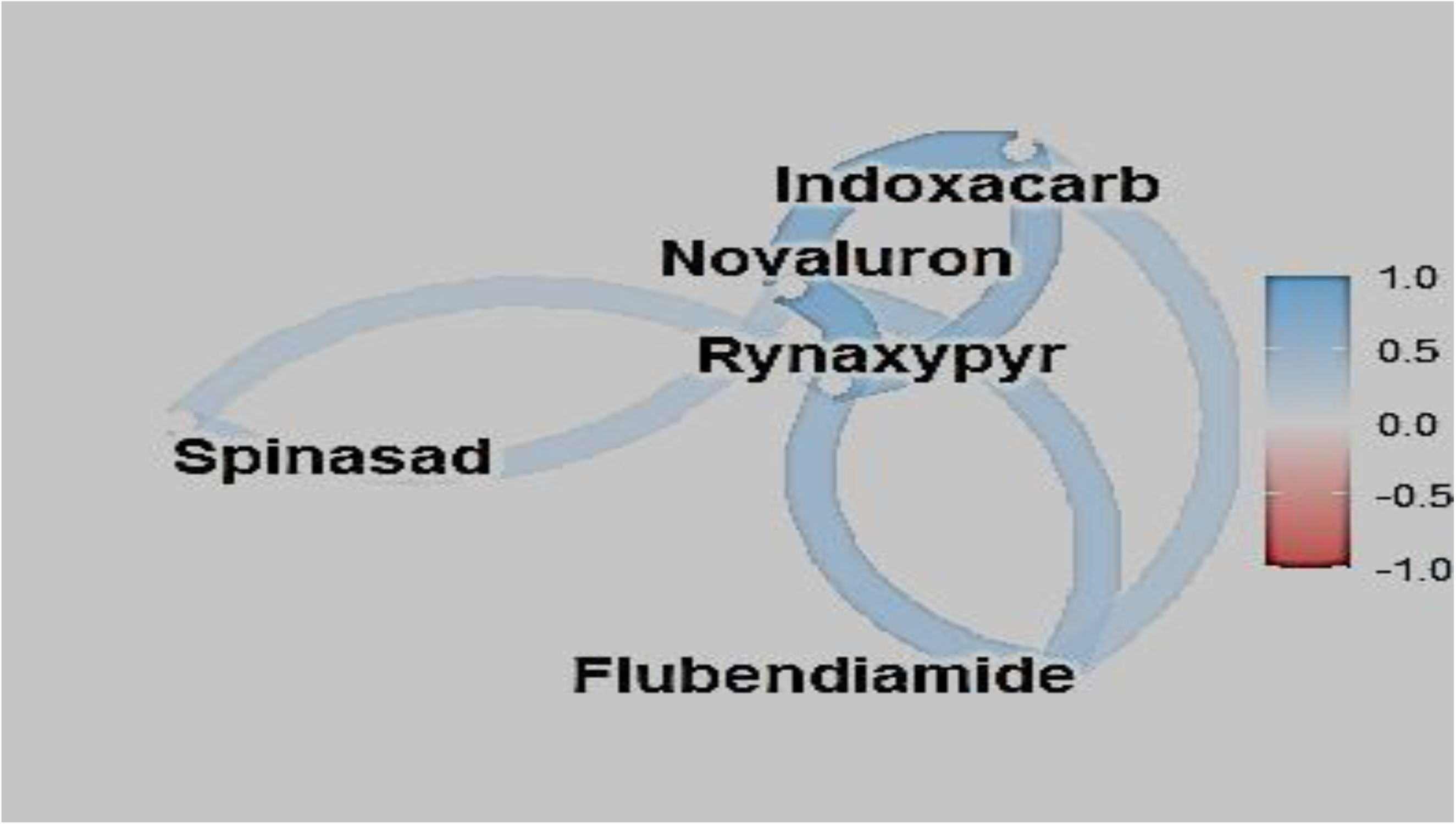
Pairwise correlation coefficient comparisons between log LC 50 values of the insecticides tested on the field populations of *Helicoverpaarmigera*.

RCH strain from Raichur location was preeminent in all the morphometric parameters larval length (2.75±0.48 cm), larval weight (0.511±0.04 g), pupal length (1.76±0.18 cm) and pupal weight (0.309±0.05 g) and positive correlation discern in both groups of insecticides with the larval weight and pupal length as well as pupal weight, alike positive and a significant correlation (0.781* and 0.681*), respectively, in the case of both insecticidal group with a larval length of *H. armigera*as well. Correspondingly Fakrudin*et al*. (2004) distinguished mean pupal length of Raichur population was found to be highest (1.75 cm) among different locations of South India which includes Karnataka into the bargain. Correlation between morphometric parameters and conventional insecticides cypermethrin resistance levels showed a positive and significant correlation with larval length and pupal weight that looks similar uniquely for larval length in the current study even so for newer and IGR insecticides.

Morphometric parameters of *H. armigera* across geographical populations of Karnataka along with susceptible lab strain Morphometric observation was seen in RCH strain from Raichur location was pre-eminence in all the morphometric parameters range and mean of larval length (2.00-3.25 cm and 2.75±0.48 cm), larval weight (0.460-0.600 g and 0.511±0.04 g), pupal length (1.50-2.00 cm and 1.76±0.18 cm) and pupal weight (0.22-0.39 g and 0.309±0.05 g) which was followed by KLB strain (Kalaburgi). Similarly, Parmar and Patel (2018) larval and pupal, lengths and weights of *H. armigera* were significantly more in case of field populations of Vadodara and Ahmedabad locations as compared to laboratory reared susceptible strain.This evident different locations strain morphological parameters might be influenced by ecological factors, habitat and feeding behaviour of *H. armigera*. Pigeonpea crop is dominant in area and production compared to other crops in Kalaburgi and Raichur regions of Karnataka, and it is a major source of nutrient for *H. armigera* it can expect more dietary requirements from the crop where the maximum resistance level tested against insecticides as well in the study.

Morphometric correlation (Table 7) with insecticidal resistance significantly positive between larval length in Novaluron (IGR), resistance development may be due to xenobiotics being responsible for better phenotypic characteristics; it is very difficult to say. Higher phenotypic attributes in terms of higher larval length might have ascended the physiology of the body with escalated enzymatic action enabling larvae to tolerate higher doses or titer of insecticide (Fakrudin*et al*., 2004). Similarly, Michael and Jane (1990) found relationships between larval weight and degree of resistance to cypermethrin in tobacco budworm, *Heliothisvirescens* (F.) indicated that the expression of tolerance and dynamics of tolerance within populations are mediated by even small-scale weight variation, that relationships between tolerance and weight depend on the dose, and it may be possible to refine resistance predictions by accounting for weight variation in treated populations which will support this study.

From the survey study, the pesticide use pattern in the case of *H. armigera* on major cropping systems (Honnakerappa and Udikeri, 2018) apart from that some aspects of GAPs (Good agricultural practices) pertaining to the resistance studied through the survey. The majority of the farmers had far below knowledge on insecticide recommendations as per the Insecticide Act, toxicity classification, and most farmers contact pesticide dealers for spraying recommendations because they are the only source of advisers in rural areas. More than 60 percent level of education in farmers is almost nil hence due to illiteracy and lack of knowledge the farmers usually go for pesticide spray more frequently with double or more than double of the recommended dosage and they fail to follow instructions given in the leaflets of any insecticide pockets or bottle, could be the reason farmers use a high amount of insecticides for spray. Increased application of insecticide for protecting crops against insect pests poses critical challenges as it may accelerate widespread resistance in strains (Ranson*et al*., 2011).

The majority of farmers were not practicing GAP’s related resistance management viz., monitoring pest level by keeping sex pheromone traps which give clue for initiate spray against pest here only 15.83 percent of farmers were monitoring the pest, 18.33 percent were practicing refugia strategy (Table 8). Earlier cotton seed companies used to give non-Bt cotton and pigeonpea seeds with *Bt* cotton to restore susceptible strain of *Bt* resistance pest populations that could break down the resistance in current pest field populations but farmers were in a delusion of growing non-Bt seeds not worth the yield. Usage of bioagent NPV/ *Bt*is one of the best components in IPM and IRM but neglect should be avoided for resistance management. Pesticide usage pattern significantly high in conventional insecticides (50 percent) and in reaming 50 percent seen in these newer group (19 %), IGRs insecticides (27 %) and 4 % in *Btk*(Fig. 4) it will clearly decipher the pattern of resistance level seen in *H. armigera*from different locations of Karnataka among different group of insecticides which were studied in Lab (Honnkaerappa and Udikeri, 2021).

## Abbreviations

HVR: Haveri
VJP: Vijayapur
DWD: Dharwad
BLG: Belagavi
SMG: Shivamoga
KLR: Kolar
RCR: Raichur
KLB: Kalburgi
GDG: Gadag
Btk: Bacillus turengensiskurstaki

## Notes

### Competing Interest Statement

The authors have declared no competing interest.

